# Autism-associated oxysterol regulates GABAergic neurogenesis and subtype fates

**DOI:** 10.1101/2025.07.01.662540

**Authors:** Maria Cruz-Santos, Ethan Kidd, Zongze Li, Daniel Cabezas De La Fuent, Sara Davies, Ngoc-Nga Vinh, Marija Fjodorova, Meng Li

**Affiliations:** Neuroscience and Mental Health Innovation Institute; Division of Psychiatry and Clinical Neuroscience, School of Medicine; School of Bioscience, Cardiff University

**Keywords:** human pluripotent stem cells (hPSCs), neuropsychiatric disorders, neurogenesis, oxysterol, liver X receptor

## Abstract

Distorted GABAergic neurodevelopment is believed to underscore cortical network dysfunction that lies at the heart of neurodevelopmental disorders (NDD) such as autism and schizophrenia. GABAergic neuron diversity is sculptured by cortical environmental cues during protracted postmitotic differentiation. However, the mechanism by which the NDD environment influences GABAergic neuronal development remains largely unknown. Oxysterols are oxidized metabolites of cholesterol that can interact with developmental signaling pathways. Using an iPSC model recapitulating human forebrain GABAergic neuron development and single-cell transcriptomic profiling, we show that 24S, 25-epoxysterol, an NDD-affected oxysterol highly enriched in the fetal brain, promotes neurogenesis, and disturbs the composition of GABAergic neuronal subtypes. Moreover, pharmacological and genetic interrogation identified the liver X receptor as a regulatory pathway mediating the action of 24S, 25-epoxysterol. These findings provide insights into the roles of cholesterol metabolism in neuronal development and a potential mechanism by which dysregulated brain oxysterols contribute to the pathogenesis of NDD.

## Introduction

Gamma-aminobutyric acid (GABA)-releasing inhibitory neurons are a highly diverse population of cells that play a critical role in maintaining neural circuit functions and modulating complex behaviors^1,2^. Dysfunction of these neurons leads to an excitatory-inhibitory imbalance that lies at the heart of neurodevelopmental disorders (NDDs), such as schizophrenia, autism spectrum disorders, and epilepsy^1,3–5^.

Disturbances in GABAergic neuron function have also been linked to altered network activity and cognitive dysfunction in Alzheimer’s disease^6^. A growing body of genomics studies has demonstrated that NDD risk variants are over-represented in GABAergic neuron transcripts and open chromatin and are already evident in fetal development^7–11^. These findings suggest an early developmental etiology of these conditions.

24S,25-epoxycholesterol (24S,25-EC) is the most prevalent oxysterol in the fetal brain and plays a critical role in midbrain dopamine neurogenesis^12–17^. We recently reported that 24S,25-EC production is altered in autism iPSC-derived cortical cells, leading to either a delay or accelerated cortical neuron differentiation, depending on the direction of the 24S,25-EC level change caused by distinct disease genetic alterations^18^. While originating from the medial ganglionic eminence (MGE), forebrain GABAergic interneurons specify into diverse subtypes in the cortex and striatum, where the local environment sculptures this process. Moreover, 24S,25-EC can activate hedgehog signaling by binding to its signal transducer smoothened^19^.

Together, these findings raise the question of whether 24S,25-EC modulates MGE neural progenitor induction and their differentiation into MGE-derived neuronal subtypes and how changes in its production in pathological brain environments disrupt this process.

Using single-cell RNA sequencing (scRNA-seq), here we mapped the transcriptional landscape of iPSC-derived MGE neurons against that of human MGE. Our analysis confirmed the in vitro production of major GABAergic neuronal subclasses with a high-fidelity match to human MGE cell populations, reinforcing the suitability of iPSC-based GABAergic neuron models for studying inhibitory neuron development and NDD pathogenesis. We report that 24S, 25-EC promote neurogenesis via signaling through LXRβ, leading to alterations in the neuronal subtype composition. This study provides novel insights into the regulatory role of 24S, 25-EC on MGE neural progenitors and a potential mechanism by which dysregulated brain oxysterols contribute to the pathogenesis of NDD.

## Results

### 24S,25-EC promotes LHX6^+^ cell production from hiPSCs without affecting ventral patterning

Given its prevalence in the fetal brain and its high-affinity binding to smoothened in vitro^19^, we first tested the ability of 24S,25-EC to ventralize forebrain neural progenitors. KOLF2 iPSCs were first induced into forebrain neural progenitors without patterning cues^20^, followed by exposure to 1mM 24S,25-EC between days 10 and 18 (d10-18, Figure S1A). A published MGE induction protocol was used as the positive control^21^, where cultures were treated with sonic hedgehog (SHH) and smoothened agonist, purmorphamine (PM), in the same time window. Efficient forebrain neural induction was achieved under all conditions, as demonstrated by the highly enriched presence of FOXG1 at d20 (Figure S1B-C). As expected, most cells in SHH-treated positive control cultures expressed the MGE progenitor markers NKX2.1 and OLIG2 (Figure S1D-E), and a small population of cells expressed the CGE marker COUTF2 (Figure S1F), while few were found in the basal control cultures. However, 24S,25-EC treatment did not lead to an increase in NKX2.1^+^ cells over the basal condition, suggesting a lack of smoothened agonist activity.

We next investigated whether 24S,25-EC could affect neuronal production from SHH patterned MGE progenitors^22,21^. To this end, 24S,25-EC was added at a later stage between d15-25 of differentiation (Figure 1A). For ease of analysis, this experiment utilized an LHX6-mEmerald reporter human iPSC line that faithfully expresses the genetic knock-in mEmerald in LHX6-expressing neurons^23^. LHX6 marks migrating MGE-derived cortical interneurons and some GABAergic projection neurons in vivo. We observed a gradual increase of mEmerald^+^ cells from d25 till d50 in both the basal control cultures and those treated with 0.5μM and 1μM 24S,25-EC (Figure 1B-C). However, there were significantly more mEmerald^+^ cells in 24S,25-EC treated cultures than in the controls at d25, d30 and d40, although the control cultures caught up by d45. In contrast to the increase in mEmerald^+^ cells, protein markers expressed by dividing MGE progenitors, such as NKX2.1, OLIG2 and FOXG1, decreased in cultures exposed to 1μM 24S,25-EC compared to the controls at d25 (Figure 1D-E). We observed no differences in the expression of COUPTF2, a marker of dorsal CGE and LGE.

**Figure 1.**
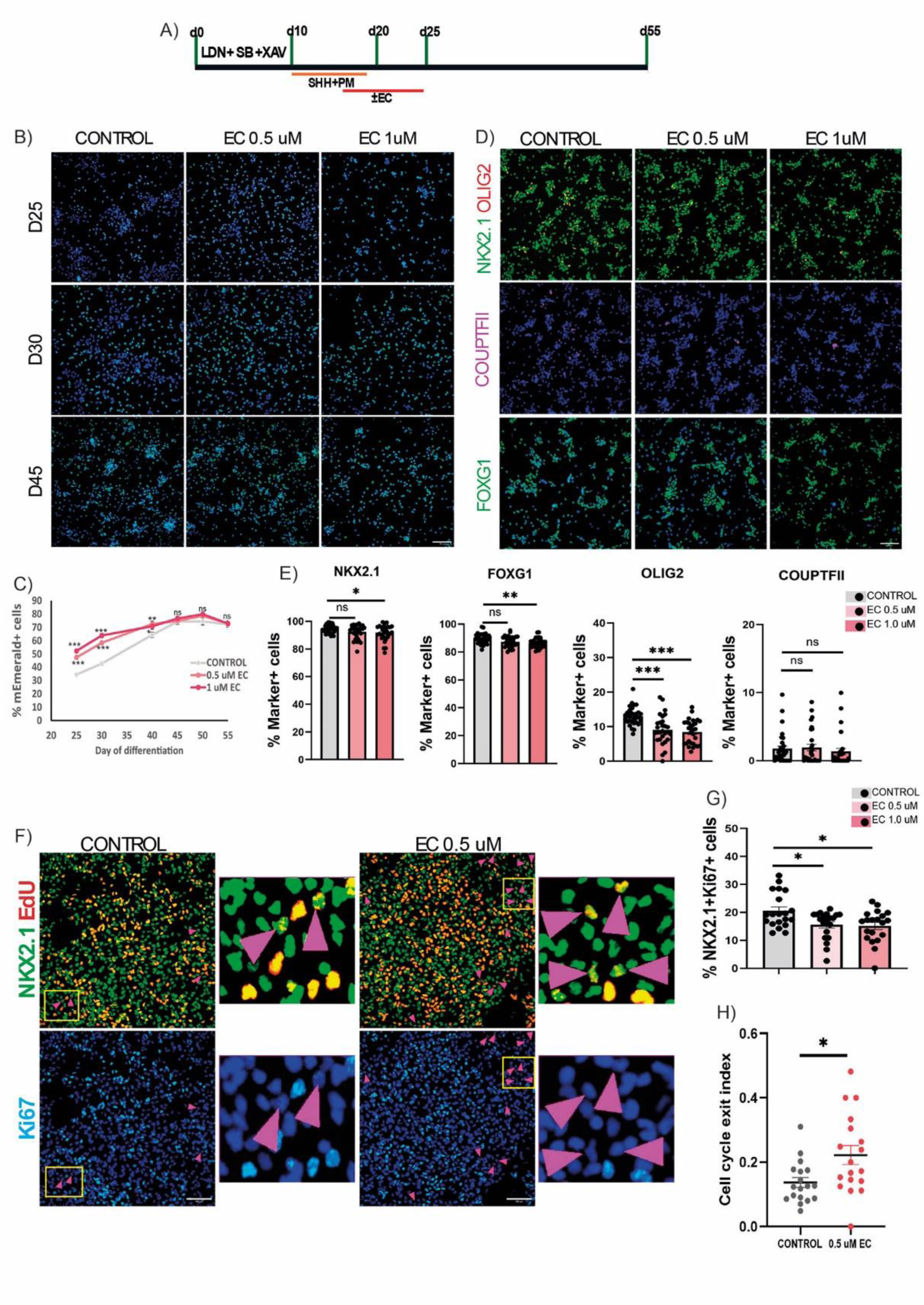
24S, 25-EC promotes neuronal production from NKX2.1^+^ progenitors. A) Outline of the neural differentiation paradigm and 24S, 25-EC treatment regime. B) Endogenous mEmerald signal from the LHX6-mEmerald cultures treated with vehicle or 24S, 25-EC (EC) at d25, d30 and d45. C) Quantification of mEmerald^+^ cells from d25-d55. D) Top and middle rows, triple immunostaining for NKX2.1(green), OLIG2 (red), and COUPTF2 (magenta) and single staining for FOXG1 (green) at d25 of differentiation. E) Quantification of NKX2.1^+^, FOXG1^+^, OLIG2^+^ and COUPTF2^+^ cells shown in D. F) Double immunostaining for NKX2.1 (green) and Ki67 (light blue) combined with EdU labelling (red). Arrow heads indicate NKX2.1^+^EdU^+^Ki67^-^ cells. G) Quantification of cycling NKX2.1^+^ Ki67^+^ progenitors. H) Quantification of cell cycle exit index. Data presented are mean ± s.e.m. of three independent differentiation runs each with 2-3 technical replicates. Each dot in the graphs represents an image counted. The numbers of technical replicates and images counted in C, E G, H are provided in table S4. EC treated vs vehicle control groups were compared by two-way ANOVA with post-hoc Bonferroni in C. One-way ANOVA with post-hoc Dunnett T3 or non-parametric Kruskal-Wallis with post-hoc Dunn in E. One-way ANOVA with post-hoc Bonferroni in G or Dunnett T3 in H (*p<0.05; **p<0.01; ***p<0.001). Nuclei were counterstained with DAPI (blue) or Hoechst for live imaging in B. Scale bars: 100 µm.

To investigate the cellular mechanism by which 24S,25-EC acts, we exposed d25 cultures to a 2-hour EdU pulse followed by triple immunostaining staining for EdU, Ki67, and NKX2.1 the next day. This experiment allowed for the determination of the cell cycle exit index, which is the ratio of EdU labelled progenitors that have already left the cell cycle (NKX2.1^+^EdU^+^Ki67^-^) to the total number of EdU^+^ cells. 24S,25-EC treatment led to an increase in the cell cycle exit index compared to that in the controls (Figure 1F-H). In contrast, we observed a decrease in cycling NKX2.1^+^ MGE progenitors in response to 24S 25-EC. Thus, 24S,25-EC promoted the cell cycle exit of MGE neural progenitors.

### High MGE specificity and close transcriptomics proximity of iPSC derivatives to human fetal MGE

To better characterize the iPSC-derived neurons and the effects of 24S,25-EC, we performed scRNAseq at two developmental stages: d25 and d40, on vehicle control and 24S,25-EC-treated cultures following the previously described treatment length. After quality control and batch effect correction with harmony (Figure S2A-B), we obtained 18,579 protein-coding genes out of 114,787 qualifying cells (58,066 control and 56,721 oxysterol-treated). Unsupervised Louvain clustering identified 11 cell clusters with each cell cluster containing different proportions of d25 and d40 samples, respectively (Figure 2A-C, Figure S2C-D). For both the control and 24S,25-EC-treated cultures, d25 samples contained more S or G2/M cells than G1 phase cells, as predicted by the CellCycleScoring function in Seurat, while the majority of d40 samples consisted mostly of G1 cells (Figure 2D-E, Figure S2E-F), confirming the temporal progression of neuron production as observed by the LHX6 reporter.

**Figure 2.**
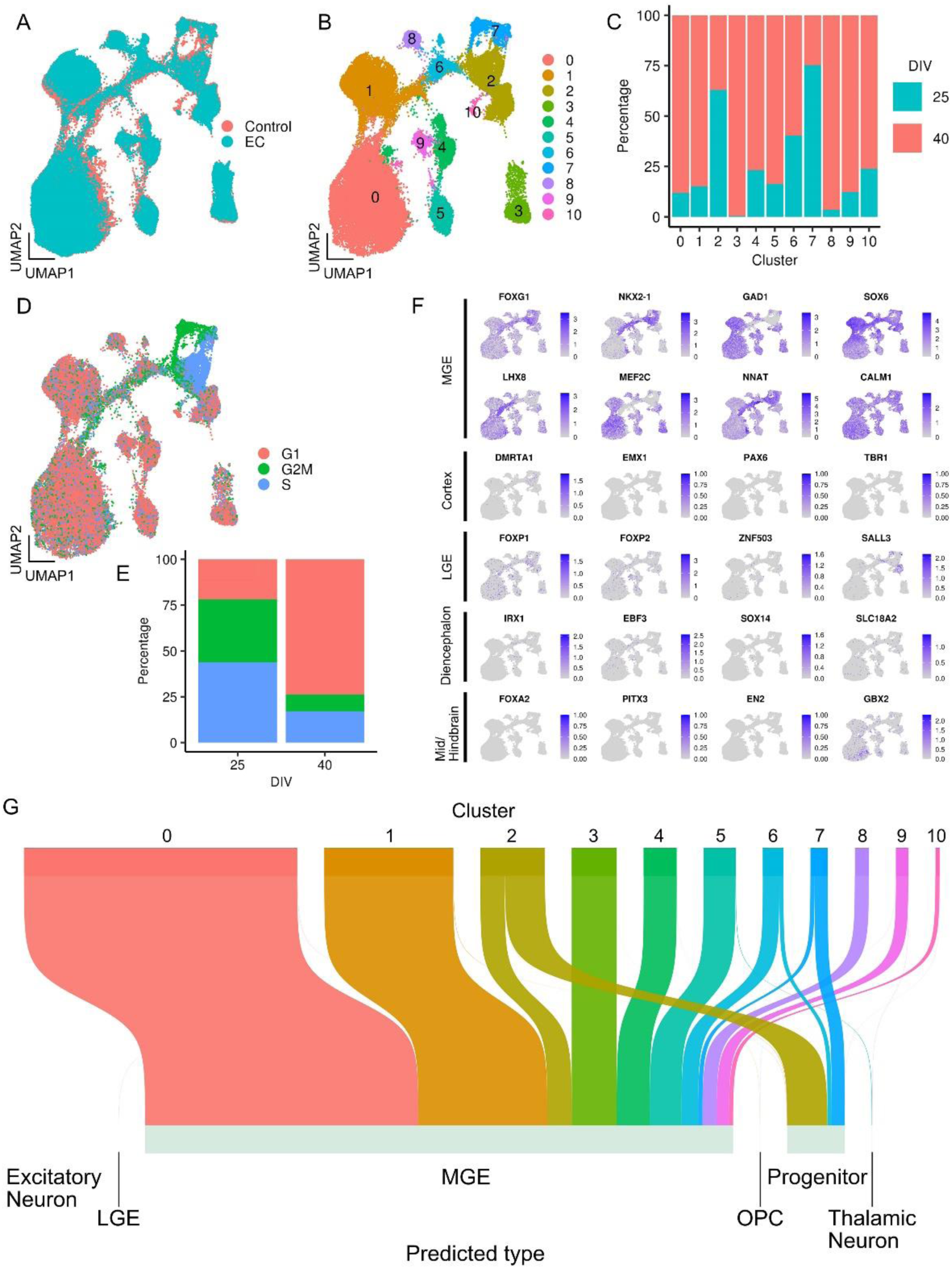
Single cell RNAseq confirms highly specific MGE fate induction A) UMAP embedding of scRNA-seq data of 58,066 control and 56,721 EC-treated iPSC-derived MGE cells at d25 and d40. B) UMAP of control cells only at d25 and d40. C) Stack plot depicting quantification of d25 and d40 in each cell cluster of the control samples. D) UMAP plot of control cells color-coded by predicted cell cycle phases. E) The proportion of control cells in each cell cycle phase at d25 and d40. F) Feature plots showing the expression levels of brain-region-specific genes. G) Sankey diagram showing alignment of iPSC-derived MGE neurons to human whole ganglionic eminence cells at 9-18 GW (Shi et al. 2021).

Consistent with the high proportions of FOXG1^+^ and NKX2.1^+^ cells on d25 (Figure 1D, E), FOXG1 and NKX2.1 transcripts were detected in all cell clusters, with higher levels concentrated in cycling cells (Figure 2D, 2F). Abundant expression of other known MGE-specific genes was also evident, including NNAT, SOX6, LHX8, MEF2C, CALM1, and the pan-GABAergic marker GAD1 (Figure 2F). NNAT was preferentially detected in S or G2/M cells, while the others were enriched in G1 dominated cell clusters, consistent with MGE neuronal gene markers (Figure 2D, 2F). In contrast, transcripts restricted to or preferentially expressed in the dorsal telencephalon (cortex, DMRTA1, EMX1, PAX6, and TBR1) or LGE (FOXP1, FOXP2, ZNF503, and SALL3) were detected in a few cells and/or at low levels (Figure 2F).

Moreover, there was no evident expression of markers indicative of other brain regions, such as the diencephalon (IRX1, EBF3, SOX14, and SLC18A2), midbrain, and hindbrain (FOXA2, PITX3, EN2, and GBX2).

We further determined the authenticity of iPSC-derived neurons by mapping our data to a published scRNAseq dataset obtained from human fetal whole ganglionic eminence using FindTransferAnchors followed by the MapQuery function of Seurat^24,25^. This analysis revealed that most iPSC clusters (0, 1, 3-5, 8-10) were mapped to fetal MGE neuron component of the human samples, although a small fraction of cluster 5 (1.27 %) and cluster 9 (0.85%) cells, representing 0.088% of total cells, were predicted as thalamic neurons (Figure 2G). Most cells in clusters 2 and 7 were mapped to human fetal neural progenitors with a small population to the human fetal MGE, whereas cells in cluster 6 were predominantly mapped to the fetal MGE neurons of Shi et al. with a small fraction of the fetal progenitors. Notably, few cells were mapped onto other neural cell types present in the human fetal dataset, such as LGE (0.01% of total cells) and oligodendrocyte progenitor cells (OPC, 0.02% of total cells) and cortical excitatory neurons (0.0008% of total cells) (Figure 2G).

These findings indicate that the hESC-derived cell types are transcriptionally aligned closely to the human fetal MGE.

Schizophrenia disease genes are enriched across iPSC-derived MGE neuron populations The analysis described above led to the identification of clusters 7, 2, and 6 as progenitors, late progenitors, and nascent neurons, respectively (Figure 3A-B), whereas clusters 0, 1, 3-5, 8-10 were assigned to neurons based on their preferential expression of neuronal genes (Figure 3B). These neuronal clusters were further annotated using known genesets representing three classes of human fetal MGE neurons previously defined by Mayer et al^26^ (Figure S2G, Figure 3C). Cluster 1 was designated as B1 SST-CCK cortical interneurons, as they showed high levels of similarity to branch 1 human fetal MGE cells exhibiting transcriptional characteristics of cortical interneurons expressing PLS3, DLX2, ARX, EPHA5, NXPH1, DCLK2, and SST (Figure 3C, Table S1). Clusters 4, 5, and 9 feature branch 2 GABAergic projection neurons that show enriched expression of RALYL, PCDH7, NRP1, ZFHX3, LRP1B, RELN, and TLE4 compared to other neuronal clusters. Cluster 4 (brach 2 group 1) neurons also show differential expression of EBF1 (Figure 3C). Clusters 0, 3, and 8 showed high transcriptomic alignment to branch 3 neurons, which were enriched with SHISA9, CNTANP2, ADARB2, ENC1, and LHX8 in comparison to branch 1 and branch 2 neurons. Descendants of this branch account for most cholinergic projection neurons, cholinergic striatal interneurons, and GABAergic globus pallidus projection neurons34,35. We found that Cluster 10 cells shared the expression of all the branch-specific genes mentioned above as well as higher levels of NKX2.1 and SFTA3 compared to other neuronal clusters; therefore, we designated these cells as unspecified neurons (Figure 3C).

**Figure 3.**
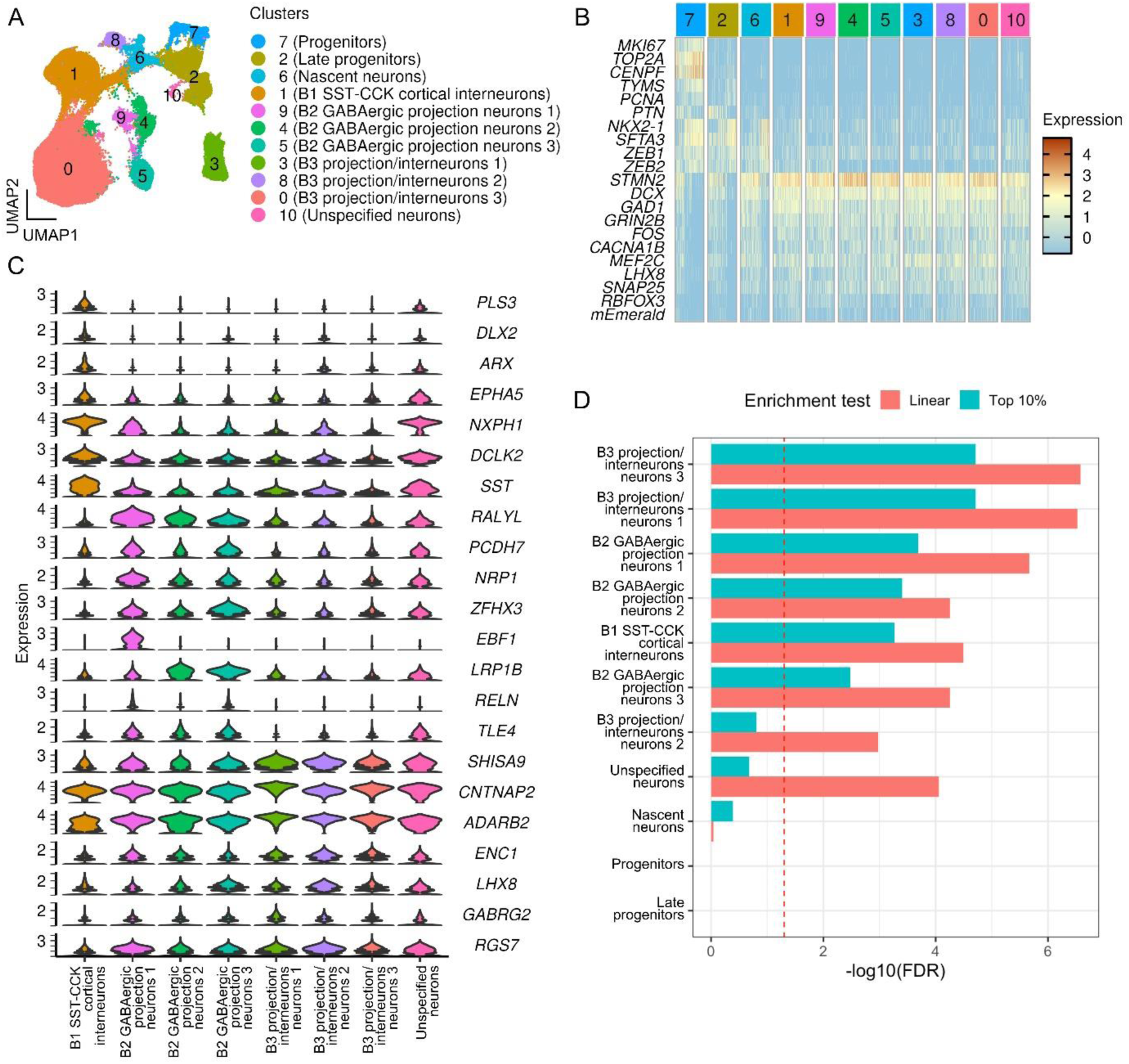
Cell-type-specific enrichment of schizophrenia GWAS signals in MGE-derived neuronal populations A) Annotation of cell types visualized by UMAP. B) Heatmap showing classic progenitor-and neuron-expressed genes in each cluster group. C) Violin plot showing differential expression of MGE neuron genes in each neuronal cluster group. D) Bar plot depicting the –log₁₀(FDR-adjusted p-values) for MAGMA cell-type enrichment tests across the annotated cell types. Two modes of enrichment analysis were tested: linear (red) using all expressed genes, and top 10% (blue) using only the top 10% most specific genes expressed in each cell type. Vertical dashed line marks the significance threshold (FDR = 0.05).

Schizophrenia risk alleles have been shown to be over-represented in human MGE-derived neurons^7,9–11^. To further assess the relevance of our iPSC-MGE cells for disease modelling, we conducted gene set enrichment analysis using the latest publicly available GWAS dataset for schizophrenia^27^. The expressed genes of all neuron groups were enriched in schizophrenia (FDR padj < 0.05, Figure 3D). To ascertain that the enrichment was specific to each cell type and not due to the genes shared by these neurons, we used the Expression Weighted Celltype Enrichment (EWCE, see method) package to calculate gene expression specificity for each group^7^. Testing only the top 10% of specific genes revealed that, with the exception of B3 projection interneuron 2, all neuronal types were still enriched for schizophrenia-associated genes (Figure 3D). Consistent with findings in the human brain, neural progenitor cells were not associated with the disease. These findings provide further support for the iPSC-MGE neurons as a suitable experimental platform for modelling neuropsychiatric and neurodevelopmental disorders.

### 24S,25-EC exhibits differential effect on neuronal subclasses

To gain insight into the effects of 24S, 25-EC on different cell populations, we compared the proportion of each cell cluster between the control and oxysterol-treated conditions. Using the whole dataset containing both d25 and d40 cells, we observed a significant reduction in cells of 24S, 25-EC treated progenitor cluster (-0.58>Log2 fold change>0.58, FDR<0.05, Figure 4A), providing independent support for 24S, 25-EC-induced cell cycle exit as measured by EdU labelling (Figure 1F-H). Intriguingly, we detected a significant reduction in unspecified neurons and an increase in two branch 3 neuronal clusters (B3-projection/interneurons 1 and 2, Figure 4A). By performing the comparison by date separately, we found that the reduction in unspecified neurons was already present at d25, whereas an increase in branch 3 neurons occurred later at d40 (Figure S2H). We also observed a reduction in the number of B1-cortical interneurons and an increase in the number of nascent neurons at d40. No significant changes in B2 neuronal clusters were detected by any means of comparison, suggesting a preferential effect of 24S, 25-EC on Branches 1 and 3 neuronal clusters.

**Figure 4.**
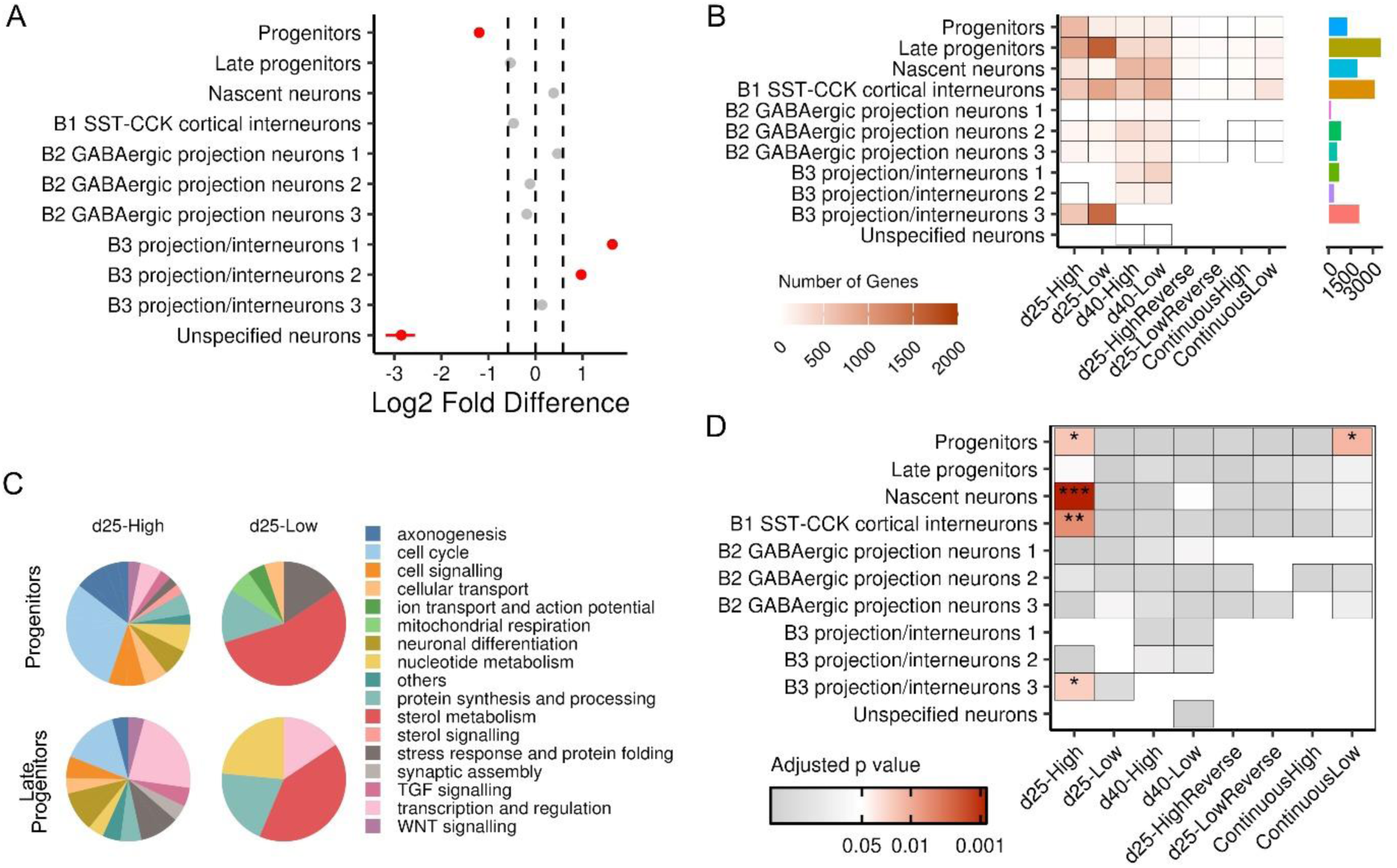
Preferential effects of 24S, 25-EC on neuronal groups A) 24S, 25-EC-dependent fold changes in the proportion of each cluster group. Significance threshold:-0.58>Log2 fold change>0.58, FDR<0.05. B) Heatmap and bar graph showing the number of DEGs of different categories by each cell cluster. C) Pie chart showing GO terms of d25 DEGs of progenitor and later progenitor populations. D) Heatmap showing enrichment of LXRβ target genes in DEGs of different categories and each cell cluster.

To explore the potential mechanisms by which 24S, 25-EC alter cellular composition, we analyzed differential gene expression between oxysterol-treated and control samples within each cell cluster considering the direction of changes at d25 and d40 in tandem. Most genes were up-or downregulated in EC-treated samples at only one time point (adjusted p value<0.05, log2 fold change ≥ 0.25). These differentially expressed genes (DEGs) were referred to as d25-High/d25-Low and d40-High/d40-Low, respectively (Table S2). A small proportion of DEGs showed the same or opposite direction of changes at d25 and d40. These DEGs are referred to as Continuous High/Low and d25-High (or low) reverse, respectively (Figure 4B and Table S2). Considering that oxysterol treatment ends at d25, we hypothesized that the d25 DEGs are likely to be enriched with direct targets and downstream pathways of oxysterol signaling, whereas the d40 DEGs may reflect secondary or long-term changes in the associated cell populations.

At d25, the progenitor and late progenitor populations responded to 24S, 25-EC with a greater number of DEGs compared to most of the neuronal clusters, as both ‘High’ and ‘Low’ (Figure 4B). B3 projection/interneuron 3 and B1 SST-CCK cortical interneurons also showed many d25 DEGs (Figure 4B). In contrast, d40 DEGs were mostly found in the nascent neurons, late progenitors, and neuronal populations (Figure 4B). These observations support the notion that 24S, 25-EC primarily acts on progenitor cells and disproportionately affects B1 SST-CCK cortical interneurons and B3 projections/interneurons among neuronal types.

However, d25-High DEGs showed contrasting GO enrichment profiles in progenitors and neurons. The progenitor and late progenitor d25-High DEGs were mainly related to the cell cycle, transcription regulation, and TGF-and WNT signalling (Figure 4C), whereas those in neurons were mainly involved in neuronal maturation and function, such as axonogenesis, synaptic assembly, and cellular transport (Table S3).

Interestingly, d25-High DEGs of progenitors, nascent neurons, B3 projection/interneurons 3 and B1 SST-CCK cortical interneurons were significantly over-represented with LXRß target genes (Figure 4D, Table S3) although B2 GABAergic neuron 2 was also enriched with targets of LXRα (Figure S3A) (https://tflink.net/)^28^, suggesting a role for LXRß in mediating oxysterol function and is consistent with LXRß being the dominant LXR isoform expressed in the brain^29^.

Moreover, as another support to EC-induced increase of neural progenitor cell cycle exit (Figure 1H), the LXRß targets identified in progenitor d25-High DEGs were mainly associated with cell cycle, transcription and regulation (Figure S3B).

Next, we performed SCENIC analysis to infer the regulatory networks underlying the actions of oxysterol. D25-High transcription factors were found to have the highest betweenness centrality measures within the cell type-specific transcriptomic networks (Figure S3C), implicating them as the likely ‘hub’ genes central to oxysterol-dependent regulatory mechanisms in the responding progenitor cells.

Crucially, many of these transcription factors, such as E2F3, HES1, TCF3/12, SMAD3, REST and YY1, are known regulators of cell cycle and neurogenesis as well as being direct targets of LXRβ (E2F3, HES1, TCF3/12, SMAD3) or its co-receptor RXR (REST, YY1)^28,30–36^.

### 24S,25-EC promotes MGE progenitor neurogenesis via LXR

Our scRNAseq analysis highlights LXR as the likely signaling receptor acting in MGE progenitors that mediates the 24S,25-EC-regulation of GABAergic neurogenesis. To provide experimental support, we pharmacologically activated LXR using the selective LXR agonist, GW3965. When applied to iPSC-MGE progenitors during d15-d25, GW3965 mimicked the effects of 24S,25-EC, as demonstrated by an increase in the number of LHX6-mEmerald^+^ neurons (Figure 5A-D). Moreover, the EdU incorporation assay showed that, similar to 24S,25-EC, GW3965 treatment resulted in a reduction in cycling NKX2.1^+^ MGE progenitors (NKX2.1^+^Ki67^+^) and an increase in cell cycle exit index (Figure 5E). In contrast, the addition of the LXR antagonist GSK2033 abolished the neurogenic promoting effect of 24S,25-EC by blocking cell cycle exit (Figure S4).

**Figure 5.**
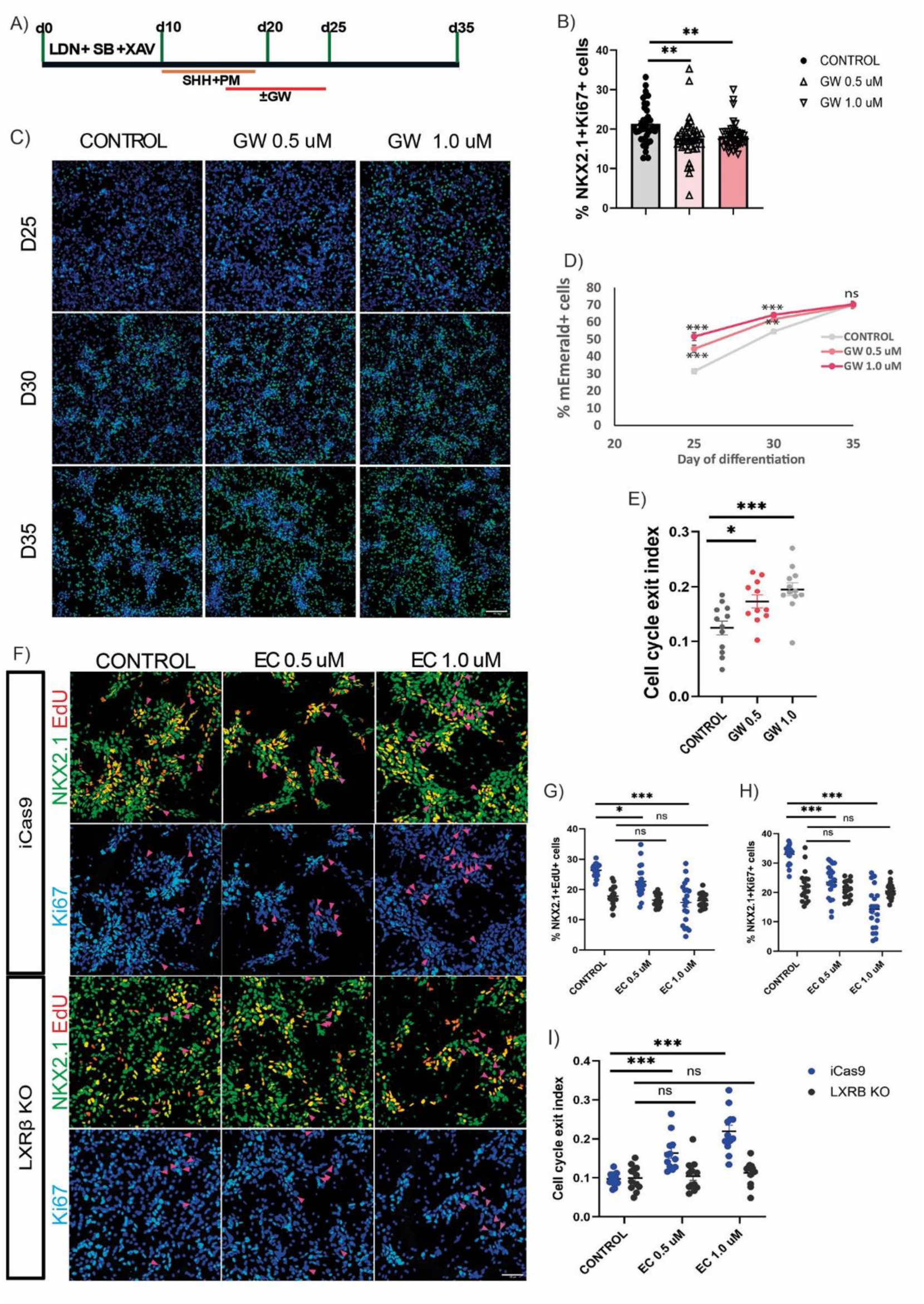
24S, 25-EC promote MGE progenitor neurogenesis via LXR A) Outline of the neural differentiation paradigm. B) Bar graph showing % of NKX2.1^+^ Ki67^+^ cells in d25 cultures treated with vehicle and LXR agonist GW3965 (GW). C) Endogenous mEmerald signal from the LHX6-mEmerald cultures treated with vehicle or GW at d25, d30 and d35. D) Line plot showing the % of mEmerald^+^ neurons at d25, d30 and d35. E) Dot plot showing cell cycle exit index of control and GW-treated cultures. F) Double immunostaining for NKX2.1 (green) and Ki67 (light blue) combined with EdU labelling (red.) Arrow heads indicate NKX2.1^+^EdU^+^Ki67^-^ cells. G-H) Dot plot showing the portion of NKX2.1^+^EdU^+^ and NKX2.1^+^Ki67^+^ progenitors, respectively, in isogenic iCas9 control and LXRβ knockout cultures in responding to EC treatment. I) Dot plot showing the quantification of cell cycle exit index in wildtype and LXRβ knockout cultures in response to EC treatment. Data presented are mean ± s.e.m. of three independent differentiation runs, each with 2-3 technical replicates. Each dot in the graphs represents an image counted. The numbers of technical replicates and images counted in B, D, E, G-I are provided in table S4. EC treated vs vehicle control groups were compared by Kruskal-Wallis with post-hoc Dunn in B. Two-way ANOVA with post-hoc Bonferroni in D. One-way ANOVA with post-hoc Bonferroni (E,I) or Dunnett T3 when equality of variances criteria was not met in G, H. (*p<0.05; **p<0.01; ***p<0.001). Nuclei were counterstained with DAPI (blue) or Hoechst for live imaging in C. Scale bars: 100 µm.

LXRβ is the predominant LXR receptor in the brain^29^. To exclude potential non-specific effects of the LXR agonist/antagonist, we re-examined 24S,25-EC on neural progenitor behavior using a CRISPR-engineered hESC line deficient in LXRβ and its isogenic wild-type control line^18^. Using the same experimental design as shown in figure 1A, we found that while 24S,25-EC treatment resulted in a decrease in S phase (NKX2.1^+^EdU^+^) and cycling (NKX2.1^+^Ki67^+^) MGE progenitors in the isogenic wild-type control culture, the same treatment had no effect on LXRβ-knockout MGE progenitors (Figure 5F-I). Together, our pharmacological and genetic approaches demonstrate that 24S,25-EC promotes neural progenitor cell cycle exit via LXRβ receptor.

## Discussion

The present study identified a new role for oxysterol-LXR signaling in cortical GABAergic neurogenesis and provided insights into a potential mechanism by which distorted oxysterol metabolism affects GABAergic neuron subtype composition that may underlie the pathogenesis of neurodevelopmental disorders.

Oxysterols are by-products of cholesterol biosynthesis and metabolism, and are capable of interacting with signaling pathways that play pivotal roles in neural development^19,37^. Altered levels of brain-derived oxysterol in the plasma or cerebrospinal fluid have been reported in individuals with neurodegenerative diseases^38–41^ and neurodevelopmental disorders^38–44^, while 24S, 25-EC was found to be produced at abnormally high levels in iPSC-derived cortical cells from autism patients and those carrying an autism risk gene mutation^18^. However, the roles of dysregulated oxysterols in disease pathogenesis remain largely unknown.

GABAergic neurons comprise an extremely heterogeneous group of cells with many specialized subtypes. Distinct neuronal subtypes populate the different layers of the adult cortex, striatum, and hypothalamus^45^. Single-cell transcriptomic studies have indicated that distinguishable GABAergic neuron subtypes emerge during fetal development, at a time when precursors migrate, morphologically mature, and establish synaptic connections to other cortical neurons^24,26,46^. Hence, changes in cortical cues will impact GABAergic neuron fates and consequently their functions.

In this study, the majority (>75%) of neurons in all neuronal clusters were born after d25, when 24S, 25-EC treatment ended. However, only three neuron populations (B3 projection/interneurons 1 and 2 and unspecified neurons) were affected, suggesting a selective effect of 24S, 25-EC either on pre-committed subtype neural progenitors or a bias on differentiation of common progenitors towards specific fates. Our regulatory network analysis of the affected neuronal populations did not reveal potential transcription factor programs that preferentially drive or inhibit the production of these neuron types but common pro-neurogenesis regulators (see below). Moreover, compared to other neuronal types, a greater proportion of B3 projection/interneurons 1 and 2 were generated after d25. Hence, there is increased chance of 24S, 25-EC effect on ‘late born’ neurons.

During development, the birthdates of MGE-derived neuronal subtypes are temporally regulated, which is a critical factor in determining the interneuron subtype diversity^47–49^. While the link between birthdates of MGE-derived neurons and interneuron subtypes has been established, the instructive mechanism that delineates how interneuron diversity is shaped in relation to the moment they exit the cell cycle is poorly understood, nor how extracellular cues might regulate this process and therefore potentially affect interneuron diversity. This study provides proof of principle that environmental changes in the developing cortex, such as altered levels of 24S, 25-EC production by cortical cells, may alter the subtype trajectory of GABAergic neurons and subsequently alter the global profile of the inhibitory-excitatory circuitry implicated in neurodevelopmental disorders.

HES1, SMAD3, and TCF3/12 were among the top regulated transcription factors identified by regulatory network analysis as likely ‘hub’ transcription factors mediating 24S, 25-EC regulation of neurogenesis (Figure S3C). As further support to this notion, all these transcription factors are direct targets of LXRβ (https://tflink.net), the presence of which is required for 24S, 25-EC-dependent neurogenesis. Interestingly, TGFβ has been shown to accelerate the generation of LHX6^+^ MGE-like neurons by promoting neural progenitor cell cycle exit^23^, while the interaction between the WNT and NOTCH pathways has been shown to play a role in regulating human MGE progenitors^50^. It is worthy of mention that WNT activation can either promote neural progenitor proliferation or differentiation depending on the context^51–56^. One such context is the presence of other signaling molecules and the crosstalk between them. Therefore, future studies are needed to experimentally delineate how LXR interacts with WNT and/or TGFβ signaling.

Importantly, our scRNAseq analysis provides unbiased evidence of the high level of transcriptomic proximity of iPSC-derived GABAergic neurons to human MGE-derived neurons in vivo, making these cells a robust platform for modelling neurodevelopmental and psychiatric disorders. Moreover, rodent MGE or iPSC-derived MGE-like neurons grafted into numerous brain regions have demonstrated their capacity to inhibit local circuitry^57–60^, encouraging the pursuit of this approach for emerging therapeutic applications. Our study indicates that directing stem cell differentiation towards a desired GABAergic subtype is possible by manipulating extrinsic signaling, hence providing grounds to tailor neuronal diversity for enhanced therapeutic outcomes.

## Materials and Methods

### HESC culture and neuronal differentiation

The hESC lines used in this study were corrected KOLF2 (KOLF2.1J) hiPSC line^61^, KOLF2-derived LHX6-emerald interneuron reporter line^23^, iCas9 and iCas9-derived LXRβ knockout line^18^. All hPSCs were cultured on Cultrex (R&D systems) coated plastics in E8 Flex media (ThermoFisher) in a humidified 37°C incubator with 5% CO_2_. Media was changed daily and cells passaged mechanically with 0.02% EDTA at 80% confluence. MGE differentiation followed procedures described previously^23^. Briefly, cells from two 80% confluent wells of a 6-well plate were plated onto a 12-well plate previously coated with Cultrex (R&D systems) in E8 media (Day 0) and changed to N2B27 the next day. Cells were induced to neuroectoderm fate by LDN-193189 (100nM, Tocris), SB-431542 (10µM, Tocris) and XAV-939 (2µM, Stratech) from day 1-10 followed by ventralisation with a combination of recombinant Sonic Hedgehog (SHH, 200 ng/ml, Peprotech) and a small molecule SHH agonist purmorphamine (PM, 1µM, Merck) from day 11-18. To promote terminal differentiation and cell survival, cultures were treated with BDNF (10ng/ml, Peprotech) from day 25 until harvest. For oxysterol experiments, 24S, 25-epoxycholesterol (0.5-1 μM, Abcam) was added to N2B27 media from day 15 to 25. For testing MGE induction effect of 24S, 25-EC (d10-18 treatment), neural differentiation follows the same procedure without SHH and PM. For pharmacological interrogation of LXR singaling, ethanol control or small molecule LXR agonist GW3965 (0.5, 1 µM, Stratech) and/or antagonist GSK2033 (0.5, 1 µM, stratech) were added during d15-25.

### Immunocytochemistry

Cultured cells were fixed in 3.7% PFA for 15 min at 4 °C. For nuclear antigen detection, an additional fixation with methanol gradient was performed, which includes 5 mins each in 33% and 66% methanol at room temperature followed by 100% methanol for 20 min at-20 °C. Cells were returned to PBS containing 0.3% Triton-X-100 (PBS-T) via inverse gradient. Cells were then permeabilized with three 10-minute washes in PBS-T and then blocked in PBS-T containing 3% BSA and 5% donkey serum. Cells were incubated with primary antibodies in blocking solution overnight at 4°C. Following three PBS-T washes, Alexa-Fluor secondary antibodies were added at 1:1000 in PBS-T for 1 hour at ambient temperature in the dark. Three PBS washes were then performed that included once with DAPI at 1:3000. Images were taken on a PerkinElmer OperaPhenix or a Leica DMI6000B inverted microscope. Quantification was carried out in Cell Profiler (cellprofiler.org) or manually using ImageJ (imagej.net) by examining 2-10 randomly selected fields of views from each of the 3 technical replicates performed in at least 3 independent differentiations (biological replicates). Each counting field is presented as a dot in relevant bargraphs with the number of counting fields for each bargraph shown in table S4. The following antibodies were used for the immunofluorescence studies: rabbit anti-FOXG1 (Abcam AB18259, 1:250), NKX2.1 (Abcam AB76013, 1:1000), OLIG2 (R&D systems AF2418, 1:200), COUPTF2 (R&D systems PP-H7147-00, 1:100), anti-Ki67 (ACK02, 1:1000), secondary antibodies (Alexa Fluor 488, 594 and 647, 1:1000)

### EdU Labelling and detection

EdU labelling and detection was performed using a Click-iT EdU Assay kit (Thermo Fisher). Cells were incubated with EdU for 2 hours at specified day of differentiation, followed by fixation in 3.7% PFA for 15 min at 4 °C. EdU detection was carried out as per manufacturer’s protocol before antibody staining. Cell cycle exit index of NKX2.1^+^ MGE progenitors was calculated as the ratio of NKX2.1^+^EdU^+^Ki67^−^ cells to the total EdU^+^ population. Image capture and quantification was performed as described in ‘Immunocytochemistry’

### Single cell RNA sequencing and data processing

For scRNA-seq, differentiating MGE cells were collected and dissociated into single cells at d25 and d40. Briefly, cells were treated with Accutase (ThermoFisher) for 5–7 minutes at 37°C. The suspension was centrifuged at 200g, 4°C for 5 minutes to remove supernatant. Cells were then re-suspended in cold PBS with 0.5% BSA and passed through a 40μm Flowmi cell strainer. Three biological replicates of the same stage were pooled together. For quality control, cell viability was assessed by trypan blue staining and confirmed to be above 80%. Cells were loaded onto the 10X Chromium Single Cell Platform and libraries were constructed using the Single Cell 3’ Library v3.1 kit (10X Genomics) as per manufacturer’s protocol. The libraries were sequenced on the Illumina Novaseq 6000 instrument using Novaseq S2 100 cycle kit aiming for a depth of approximately 150,000 reads/cell. Sequencing was run as paired-end, 100 cycles in a standard workflow.

Raw reads were processed using Cell Ranger 7.1.0 (10x Genomics) and aligned to *Homo sapiens* GRCh38-2020-A. All downstream analysis was performed using the R package Seurat (v 5.2)^62^. For quality control, a Median Absolute Deviation (MAD) method was applied using the isOutlier function from the scuttle package (1.8.4)^63^.

Cells with nFeature 2 MADs lower than the median, median nCouns above or lower than 2 MADs, and the fraction of mitochondrial genes above 2 MADs were discarded. The normalisation and variance stabilisation of molecular count data were performed using the sctransform function in Seurat^64^. This function was used to also remove confounding sources of variation such as mitochondrial mapping percentage. The batch effect was removed using harmony (v1.1.0)^65^. Uniform manifold approximation and projection (UMAP) and unsupervised Louvain clustering were performed using RunUMAP, FindNeighbors and FindClusters function based on the first 30 principal components. Cell cycle phase scores were calculated using the CellCycleScoring function based on the expression of canonical G2/M and S phase markers^66^. For annotation, cluster-specific gene markers were found with FindConservedMarkers function with min.pct=0.25, logfc.threshold=0.25, statistically significant markers p.adjust<0.05 were tested (hypergeometric test) for gene enrichment in specific makers of MGE branches previously reported in Mayer, et al. 2018^26^.

To compare the transcriptome similarity with published human GE scRNAseq data^24^, we performed anchor-based label transfer implemented in Seurat. The anchors between the query and reference datasets were identified and filtered using the FindTransferAnchors() function with the top 50 dimensions. Then, the MapQuery() wrapper function was used to project our data onto human brain data to visualise cell type similarities between datasets. Differential gene expression analyses were performed using the FindAllMarkers and FindMarkers function with Wilcoxon signed-rank test and Bonferroni correction for total number of genes included in the analysis. Proportion change analysis was performed using the scProportionTest 0.0.0.9 package^67^ with 10000 permutations followed by FDR correction. Gene ontology (GO) enrichment was performed using the enrichGO function in the clusterProfiler 4.12.6 package^68^ based on *Homo sapiens* GO term database retrieved by the org.Hs.eg.db 3.19.1 package. Gene set over-representation tests were performed using the enricher function in the clusterProfiler package. LXRα and LXRß targets were obtained from TFLink gateway (https://tflink.net), including only TF-target interactions from Chromatin immunoprecipitation and/or electrophoretic mobility shift assay experiments. SCENIC analysis was performed using pySCENIC 0.12.1^69,70^. cisTarget databases used for SCENIC analysis were the Homo sapiens - hg38 – refseq_r80_v10-SCENIC+ databases downloaded from https://resources.aertslab.org/cistarget/. SCENIC regulon enrichment score (AUC) was used for differential testing using the Wilcoxon signed-rank test with Bonferroni correction for the total number of regulons detected. Gene networks were visualised and analysed with Analyze Network in Cytoscape 3.10.2^71^

### Gene enrichment analysis

Public summary statistics associated to the latest schizophrenia GWAS^27^ were downloaded from https://www.med.unc.edu/pgc/download-results/scz/. Summary statistics were QC and cleaned using R package MungeSumstats^72^. SNPs were mapped to genes with annotation windows of 35 kb and 10 kb upstream and downstream, respectively. Dupplicated rsIDs for the same SNP were excluded from analysis. The Seurat object containing the expression dataset was converted into a CellTypeDataset object using the EWCE R package, as reported by Skene et al^7^.

Main cell type association analysis was carried out using the MAGMA.Celltyping running MAGMA v1.10^7,73^ using a combination of linear and the top 10% most specific genes for each cell type.

## Statistical analysis

RNA sequencing data were analyzed separately as described above. The SPSS Statistics 27 software (IBM) was used for all other statistical analyses. All quantified data were plotted in Prism 10 (GraphPad Software) and are reported as the mean ± SEM from at least three independent experiments, unless otherwise specified. Data were tested for normality using the Shapiro-Wilk test and for equal variance with Levene’s test before performing statistical analyses by one-way ANOVA or two-way ANOVA (depending on the number of dependent variables) with post-hoc Bonferroni for equal variances or Dunnett’s T3 test for multiple comparisons where equal variances criteria were not met. The Kruskal-Wallis test with post-hoc Dunn’s test for pairwise comparison was used when the parametric test was not suitable. The results were considered statistically significant at p<0.05.

## Acknowledgements

We thank Dr. Francesco Bedogni, Prof. Kathryn Peall and members of the M.L. laboratory for helpful discussions during this study. RNA sequencing analysis was performed using the computational facilities of the Advanced Research Computing@Cardiff (ARCCA) Division, Cardiff University. This research was funded by the U.K. Medical Research Council (grant number MR/L020807) and EU Horizon 2020 Program ‘NSC-reconstruct’ (grant number No. 874758) to M.L.

## Author Contribution

M.C-S. M.L. conceived of the study and designed the experiments. M. C-S. and E.K. carried out and analyzed the hESC and hiPSC experiments. M.C-S. S.D. and N.N.V captured the scRNA-seq experiment samples, while S. D. made the libraries. M.C-S. Z. L. and E.K. performed data analysis. D.C.F. performed gene enrichment analysis. M.F. provided guidance on scRNA-seq data analysis to M.C-S and E.K.

M.L. wrote the paper. All authors edited and approved the manuscript.

## Inclusion and diversity

We support inclusive, diverse, and equitable conduct of research.

## Supplemental information

4 supplemental figures and 5 supplemental tables are available. Table S1: Genes preferentially expressed in each cell cluster.

Table S2: Oxysterol dependent DEGs in each cell clusters, which are catergorised as d25-High & low, d40-high low, continuous high & low, d25 high-reverse, and d25 low reverse.

Table S3, GO terms associated with DEGs.

Table S4: Information on replicates and counting fields associated with all bargraphs. Table S5: Genes used for top 10% enrichment analysis.

---

**Figure S1.**
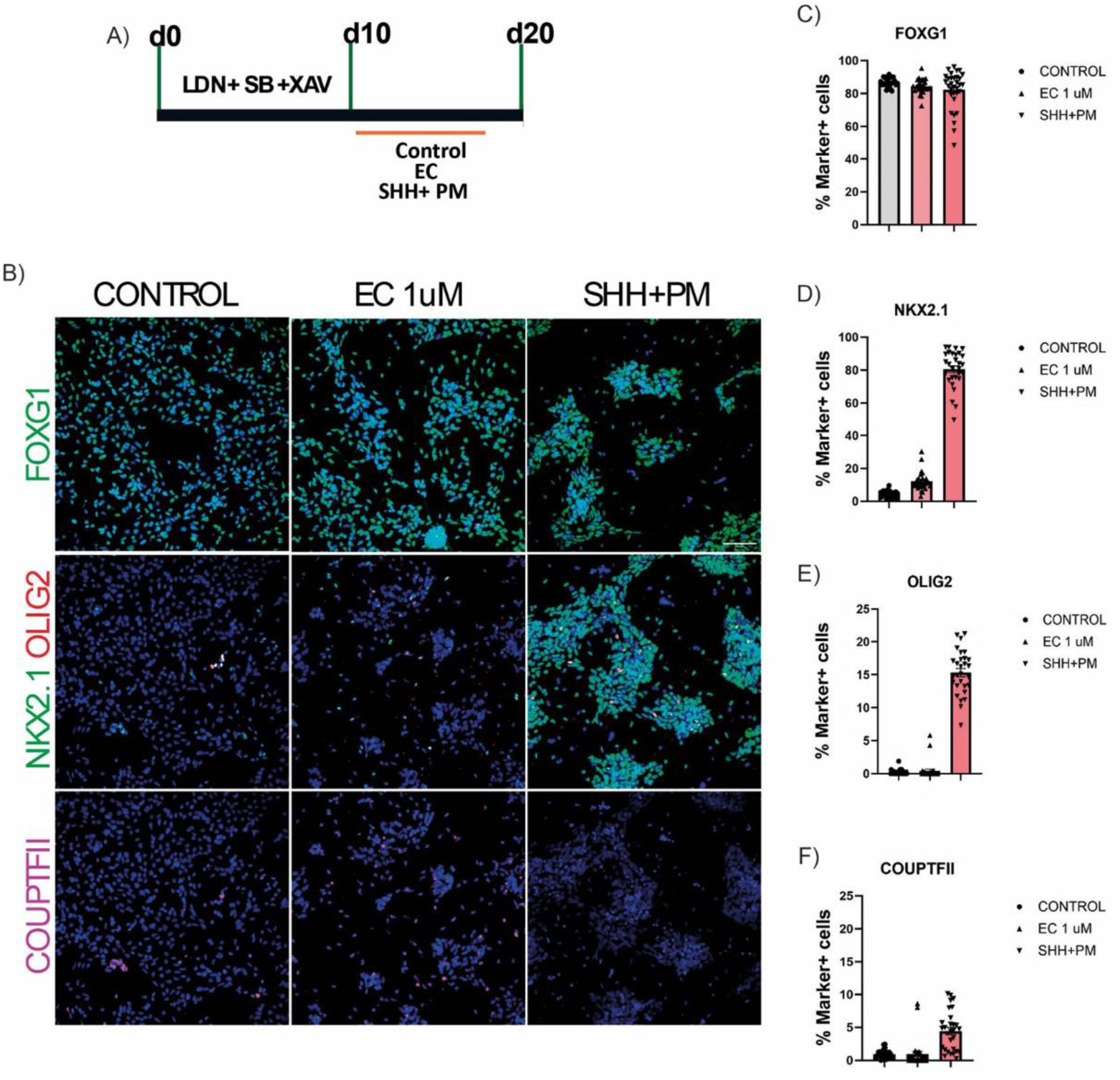
24S, 25-EC does not exhibit ventral patterning activity on PSC-derived forebrain neural progenitors. A) Outline of the neural differentiation paradigm. B) Single immunostaining for FOXG1 (green) top row and triple immunostaining for NKX2.1(green), OLIG2 (red), and COUPTF2 (magenta) middle and bottom rows at d20 of differentiation. C-F) Quantification of FOXG1^+^, NKX2.1^+^ OLIG2^+^ and COUPTF2^+^ cells shown in B. Data presented are mean ± s.e.m. of three biological replicates as described in the method section. Nuclei were counterstained with DAPI (blue). Scale bars: 100 µm.

**Figure S2.**
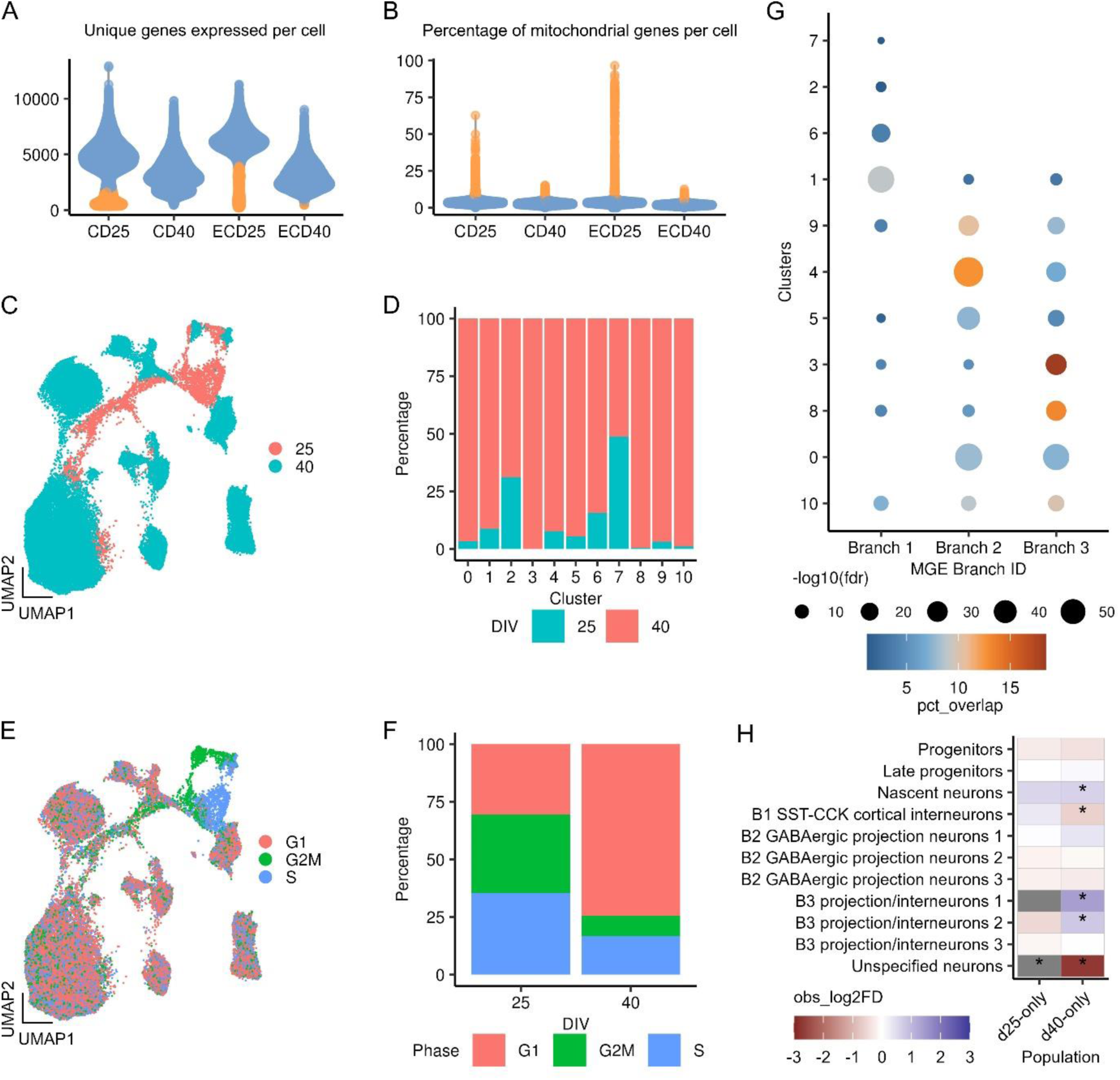
Transcriptional profiling of iPSC-derived MGE neurons, related to Figure 2 A-B) Violin plots showing the number of genes and percentage of mitochondrial genes expressed in each sample, respectively. C) UMAP embedding of 56,721 EC-treated MGE cells at d25 and d40. D) Stack plot depicting quantification of d25 and d40 in each cell cluster of the EC-treated samples. E) UMAP plot of EC-treated cells color-coded by predicted cell cycle phases. F) The proportion of EC-treated cells in each cell cycle phase at d25 and d40. G) Summary of the enrichment analysis showing transcriptional alignment of each cluster population to the three branches of human MGE cells described in Mayer et al. Dot color corresponds to percental overlap of genes, dot size corresponds to log_10_ false discovery rate. H) Heatmap showing effects of 24S, 25-EC on the proportion of each cell cluster analyzed on d25 and d40 dataset separated. Data is presented as log2 fold change. *P<0.05.

**Figure S3.**
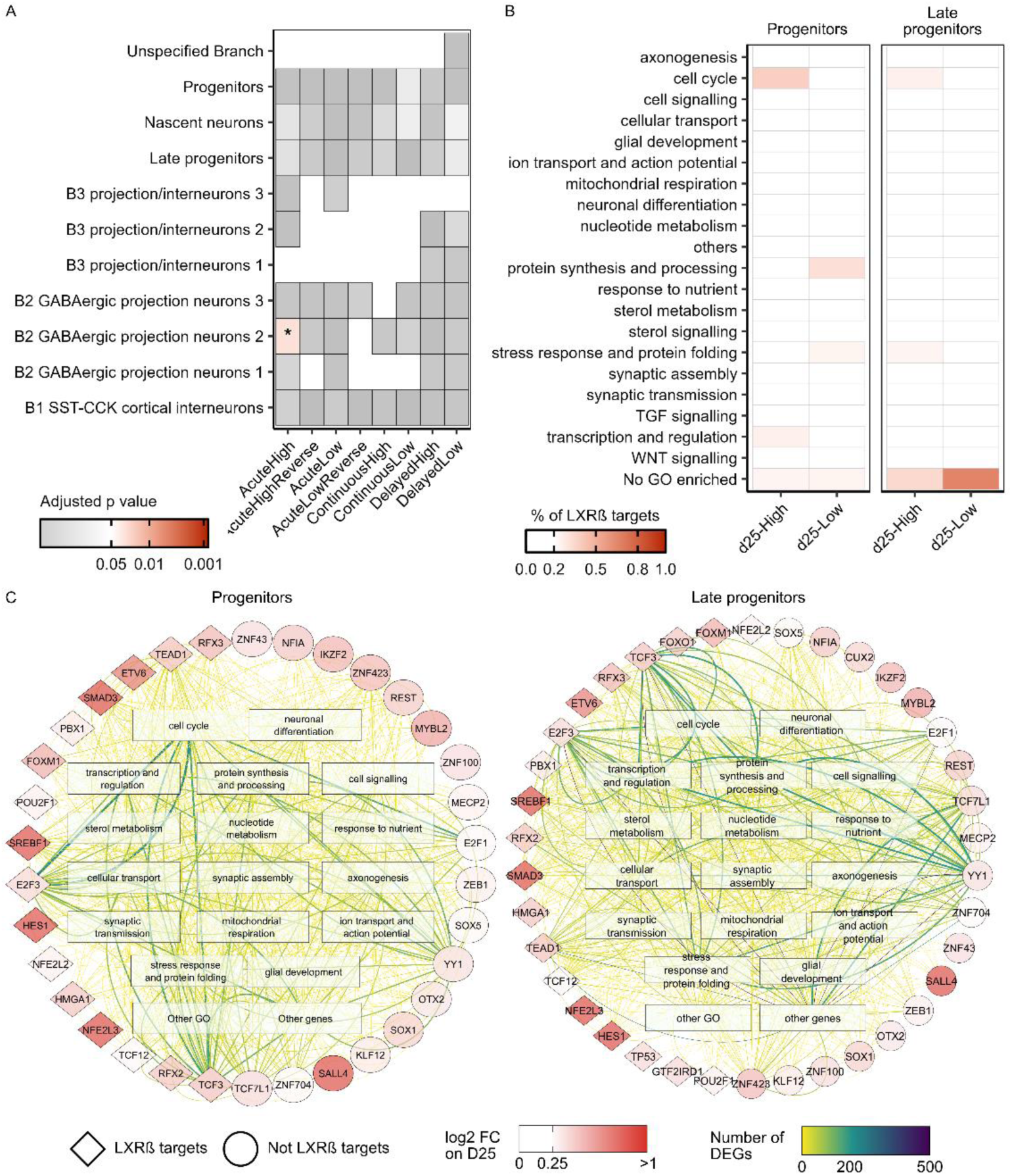
Transcriptional profiling of iPSC-derived MGE neurons, relates to figure 4 A) Heatmap showing the adjusted p values of the over-representation (hypergeometric test) of LXRα target genes in different DEG categories in each cell type. B) Heatmap showing the proportion of LXRα target genes associated with different biological processes. C) SCENIC regulatory network of d25-High transcription factors and biological process terms associated with DEGs in progenitors and late progenitors. All DEGs were represented by biological process terms. The shape of nodes depicts whether the transcription factor is a target of LXRß, while the colour represents the log2 fold-change (FC) on d25. The colour of edges represents the number of DEGs associated with the biological process terms. (GO: gene ontology; FC: fold change; DEGs: differentially expressed genes; *: p<0.05)

**Figure S4.**
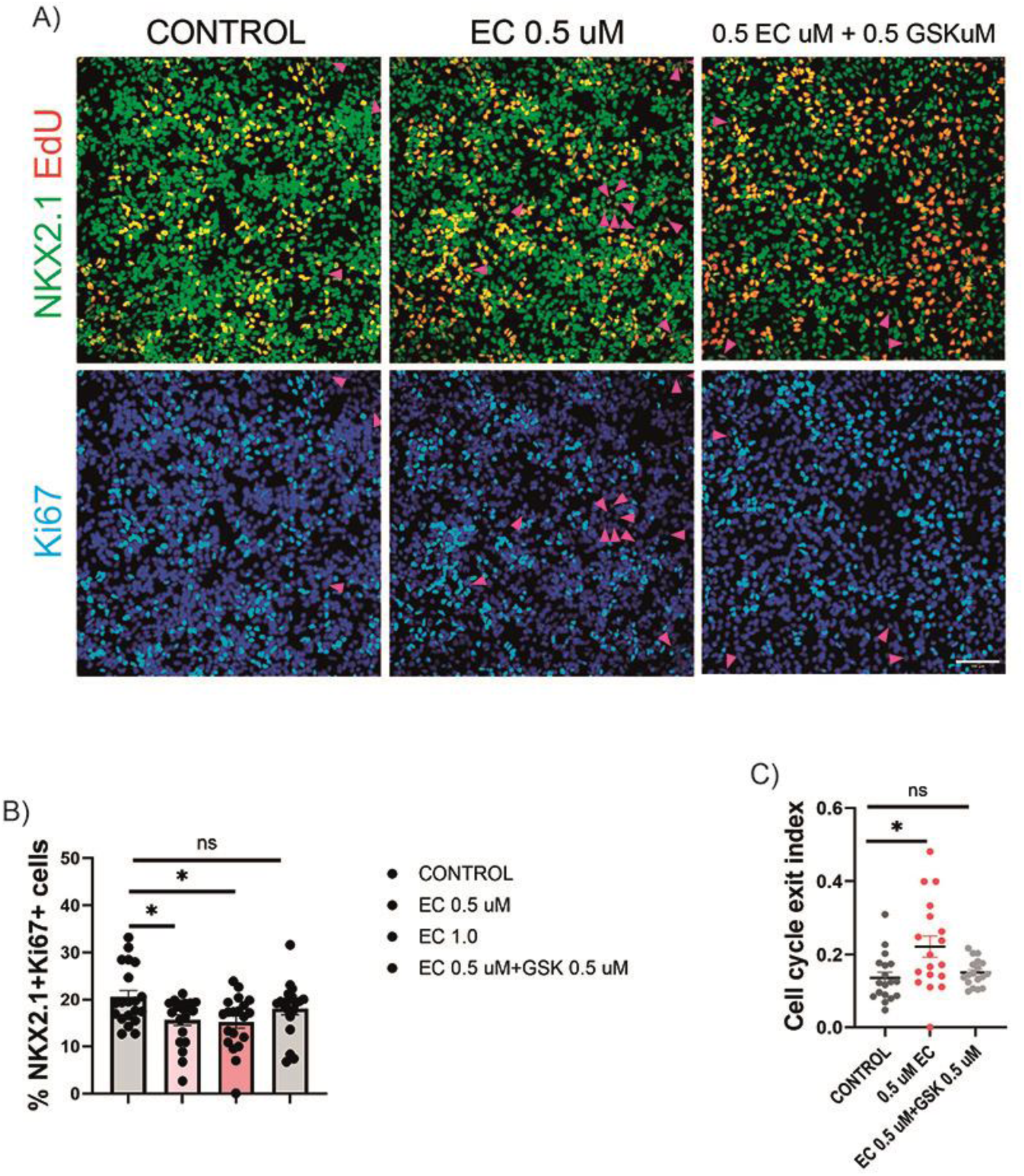
24S, 25-EC promote MGE progenitor neurogenesis via LXR, relates to figure 5 A) Double immunostaining for NKX2.1 (green) and Ki67 (light blue) combined with EdU labelling (red). Arrow heads indicate NKX2.1^+^EdU^+^Ki67^-^ cells. B) Quantification of cycling NKX2.1^+^ progenitors. C) Quantification of cell cycle exit index. Data presented are mean ± s.e.m. of three independent differentiation runs, EC treated vs vehicle control groups were compared by one-way ANOVA with post-hoc Bonferroni in B or Dunnett T3 in C (*p<0.05; **p<0.01; ***p<0.001). Nuclei were counterstained with DAPI (blue). Scale bars: 100 µm.

